# Domain-general and domain-specific neural mechanisms underlie metacognition in social and perceptual decisions

**DOI:** 10.64898/2026.11.27.769730

**Authors:** Rebekka S. Mattes, Gizem Vural, Sarina Drexler, Hongmei Xia, Ophelia Deroy, Alexander Soutschek

**Affiliations:** Department of Psychiatry, Psychosomatics, and Psychotherapy, University Hospital Würzburg, Würzburg, Germany; Department of Psychology, Ludwig-Maximilians-Universität (LMU), Munich, Germany; Graduate School of Systemic Neurosciences (GSN), Munich, Germany; Department of Psychiatry and Psychotherapy, LMU University Hospital, Munich, Germany; Comparative Psychology, Institute of Experimental Psychology, Heinrich-Heine University, Düsseldorf, Germany; Faculty of Philosophy, Philosophy of Science and the Study of Religion, Ludwig-Maximilians-Universität (LMU), Munich, Germany; Munich Centre for Neurosciences, Brain & Mind, Munich, Germany; Institute of Philosophy, School of Advanced Study, University of London, London, UK

**Keywords:** social metacognition, perceptual metacognition, domain specificity, metacognitive noise, fMRI, tACS

## Abstract

Metacognition, the monitoring and assessment of our own cognitive processes, plays a key role in our everyday decision-making across a multitude of domains. Despite this, the question of whether metacognition is implemented by domain-general or domain-specific neural processes has not yet been conclusively answered. Theoretical two-layer accounts propose that a shared core network of metacognition, consisting of prefrontal areas, might be functionally coupled with task-relevant brain regions in a domain-specific way. To test this empirically, the present study comprises three experiments. In the first experiment, participants underwent neuroimaging to compare the neural correlates of metacognitive judgements in a novel social decision task with those in an established perceptual decision task. Prefrontal areas, such as FPC, dmPFC, and dlPFC, showed domain-general activations associated with confidence, while connectivity of the FPC with task-related regions (TPJ vs. occipital cortex) was correlated with metacognition in a domain-specific way. In two further experiments, participants performed the same two tasks, while tACS was applied to selectively modulate connectivity between FPC and the TPJ or the occipital cortex, aiming to test whether this connectivity causally underlies metacognition. Indeed, the results suggest that altering FPC–TPJ vs. FPC–occipital cortex connectivity had domain-specific effects on metacognition. Together, our findings provide evidence for both domain-specific and domain-general neural mechanisms of metacognition, aligning with theoretical two-layer accounts, and advancing our understanding of the nature of metacognitive processes in different domains.

## Introduction

Metacognition describes the ability to monitor and evaluate our own cognitive performance (1), playing a central role in guiding everyday behaviour. For example, before cycling through an intersection, we assess not only whether the light is green, but also how confident we are in this perception. Metacognitive accuracy in our decisions varies across individuals, but it can also be specifically impaired in psychiatric disorders such as obsessive-compulsive disorder (2), or substance addiction (3,4). Elucidating its neural bases may therefore improve our understanding of differences and deficits and inspire training and interventions.

Neuroimaging and brain stimulation studies have associated metacognitive processes predominantly with prefrontal areas (see 5), particularly frontopolar cortex (FPC; 6,7), dorsomedial prefrontal cortex (dmPFC; 8) and dorsolateral prefrontal cortex (dlPFC; 9), which may support accurate metacognitive judgements (5,10). These regions require access to task-specific information, and FPC indeed appears functionally coupled with regions supporting first-order task processing (6,11). This raises the question of whether such connectivity differs across cognitive domains. As a second-order process, metacognition is relevant to many domains, including unisensory and multisensory perception (6,12–14), memory (15), numerical cognition (16), and value-based decisions (10), but whether it relies on domain-specific or domain-general mechanisms remains unresolved.

Behavioural evidence here is inconclusive: studies comparing metacognitive performance across perceptual modalities found moderate-to-strong correlations (17,18), whereas studies examining perceptual and other domains (e.g., metamemory) reported either moderate correlations (19–22), or none (23–25). Neuroimaging evidence, in turn, suggests a co-existence of common and dissociable brain areas implementing metacognition across domains (26–28).

The two-layer account of metacognition (29,30) may help integrate these findings: prefrontal areas like FPC, dmPFC, and dlPFC are proposed to form a domain-general, upper-layer core network that interacts with lower-layer domain-specific networks providing task-specific information for confidence judgements. This account predicts that the prefrontal core network, particularly FPC, interacts with regions involved in first-order task processing, and indeed, recent findings suggest that for several sensory tasks, confidence is implicitly encoded by relevant populations of sensory neurons (13,31). However, as outlined in a recent review (32), it remains unclear whether metacognition across and beyond sensory domains relates to, or is causally implemented by, such interactions.

Extending this framework beyond domains such as meta-perception and meta-memory generates predictions about how the prefrontal core network interacts with domain-specific regions relevant for less-studied metacognitive domains. One such domain is social metacognition, e.g., evaluating social processes like predicting others’ behaviour, which is thought to rely on perspective-taking (33), a process that has been well-studied. While perspective-taking is linked to the neural mentalising network, including the temporo-parietal junction (TPJ; 34–36), the neural basis of metacognitive knowledge about predicting others’ behaviour remains unknown. Based on two-layer accounts, we hypothesized that, first, social metacognition would be associated with activity in the domain-general prefrontal core network, and domain-specific functional coupling between prefrontal and mentalising regions such as the TPJ. Second, when comparing the neural basis of metacognition in the social domain with a more established control domain like perception, we predicted that network interactions causally underlying metacognition differ across the two domains.

To test these predictions, we conducted three experiments. First, we used neuroimaging to compare neural correlates of metacognitive judgements in a novel social decision task and an established perceptual decision task. In two further experiments, we applied bifocal transcranial alternating current stimulation (tACS) to determine whether metacognition is causally implemented by functional coupling between the prefrontal core network and domain-specific areas for first-order task performance. We selectively modulated connectivity between FPC and TPJ (experiment 2) or occipital cortex (OC; experiment 3), areas involved in social and perceptual processing, respectively. This allowed us to test whether dissociable connectivity patterns causally underlie metacognition across domains. Together, our findings provide evidence for both domain-specific and domain-general neural mechanisms of metacognition, advancing our understanding of metacognitive processes across domains.

## Results

### Behavioural task validation

To determine the neural bases of metacognition in the social domain and compare them with those underlying perceptual metacognition, we developed a novel social decision task where participants predicted whether another person would accept or reject monetary offers in an Ultimatum game (Fig 1). Such predictions of others’ behaviour are thought to rely on mentalising processes. After each decision, participants indicated their subjective confidence in the accuracy of their prediction, before receiving information on whether the other person indeed accepted or rejected the monetary offer (based on the choices of real human others). In the context of the current study, higher social metacognitive skills are indicated by increased self-reported confidence for correct versus incorrect predictions. As a non-social control task, we used an established perceptual decision task where participants rated their confidence in decisions about which of two dot clouds contained more dots.

**Fig 1.**
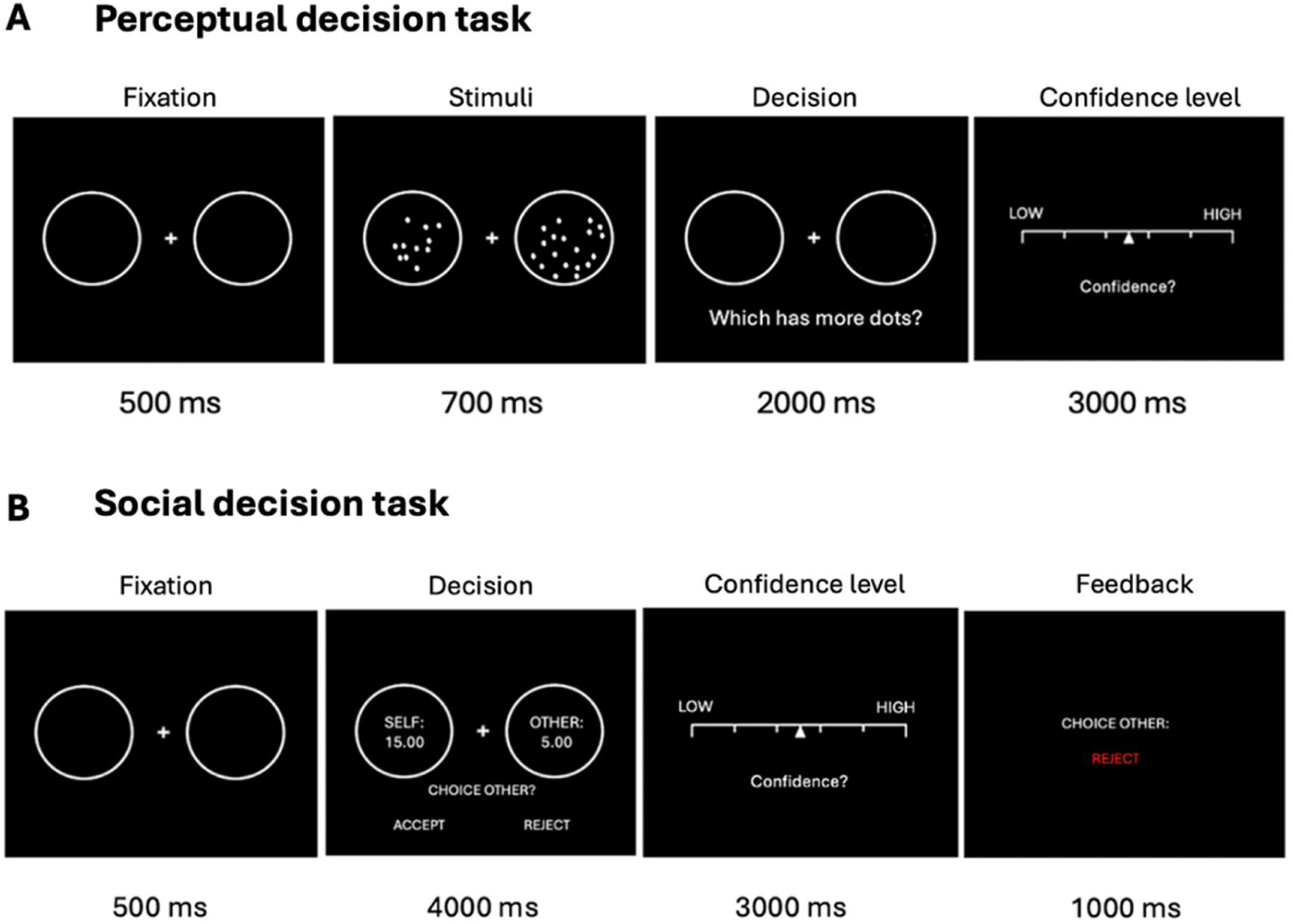
Schematic illustration of the trial sequence in both tasks. In the perceptual decision task (A), retrieved and adapted from Fleming et al. (52), participants view two circles containing various numbers of dots on a screen and decide which circle contains more dots. After each decision, they are asked to rate their confidence in their decision being correct. In the social decision task (B), the participants’ task is to predict whether other people in the role of the Responder would accept or reject the monetary offer based on the Ultimatum Game (54). Following each decision, participants are asked to report their subjective confidence that their prediction is correct. Afterwards, they receive feedback on a given person’s decision from a previous experiment.

We first assessed whether, on the behavioural level, participants possessed metacognitive access to the accuracy of their social and perceptual decisions. In a behavioural pilot experiment (without neuroimaging), we observed that participants displayed above-chance accuracy levels in both the social, ß = 0.50, *z* = 8.42, *p* < 0.001, and the perceptual decision task, ß = 1.22, *z* = 13.93, *p* < 0.001. First-level task accuracy was significantly lower for the social (mean = 62% correct responses) than for the perceptual decision task (mean = 77% correct responses), main effect of Task: ß = –0.71, *z* = 6.63, *p* < 0.001. Post-decision confidence ratings as a measure of metacognitive bias did not differ significantly between social (mean = 4.23) and perceptual decisions (mean = 3.91), as indicated by the lack of a significant main effect of Task, *p* > 0.10. Moreover, we found a significantly positive main effect of Accuracy on confidence, ß = 0.31, *t*(18) = 9.21, *p* < 0.001, suggesting that in both tasks, participants possessed metacognitive access to the accuracy of their mental performance (metacognitive sensitivity). Together, the pilot data suggest that these tasks allowed us to measure and compare metacognitive processes in the social and perceptual domains.

We replicated this result pattern in the functional imaging study, where participants performed both tasks inside the MRI scanner. Again, first-order performance accuracy was above-chance level in both tasks (social, ß = 0.53, *z* = 14.95, *p* < 0.001; perceptual, ß = 1.61, *z* = 14.19, *p* < 0.001), while significantly lower in the social (mean = 63% correct responses) than in the perceptual decision task (mean = 82% correct responses), main effect of Task: ß = –1.08, *z* = 8.98, *p* < 0.001. In the main study, we observed significantly higher confidence in social decisions (mean = 4.15) than in perceptual decisions (mean = 3.92), ß = 0.23, *t*(34) = 2.37, *p* = 0.02. Participants again possessed metacognitive access to the accuracy of their decisions (metacognitive sensitivity), as displayed through a positive main effect of Accuracy on confidence in both the social decision task, ß = 0.55, *t*(34) = 10.08, *p* < 0.001, and in the perceptual decision task, ß = 1.29, *t*(32) = 13.37, *p* < 0.001 (Fig 2A). This was robust when we controlled for Run Number to account for potential learning effects, both *p* < 0.001. Lastly, we tested whether metacognitive processes in the social and the perceptual decision tasks share common variance or are independent of each other. There was no evidence for a significant correlation in first-order task performance, *r* = –0.06, *p* = 0.72, whereas we found moderate correlations between confidence ratings (metacognitive bias), *r* = 0.51, *p* < 0.002 (Fig 2B). Interestingly, metacognitive sensitivity (confidence_correct_ – confidence_incorrect_) was not correlated between tasks, *r* = –0.09, *p* = 0.60 (Fig 2C), and neither was metacognitive efficiency (Mratio), *r* = 0.09, *p* = 0.58, nor metacognitive noise, *r* = 0.15, *p* = 0.38. While the significance of the correlations between confidence ratings may indicate that social and perceptual metacognitive judgements rely at least partially on common processes, the lack of correlations in metacognitive sensitivity, efficiency, and noise suggests that they may also possess unique, domain-specific components.

**Fig 2.**
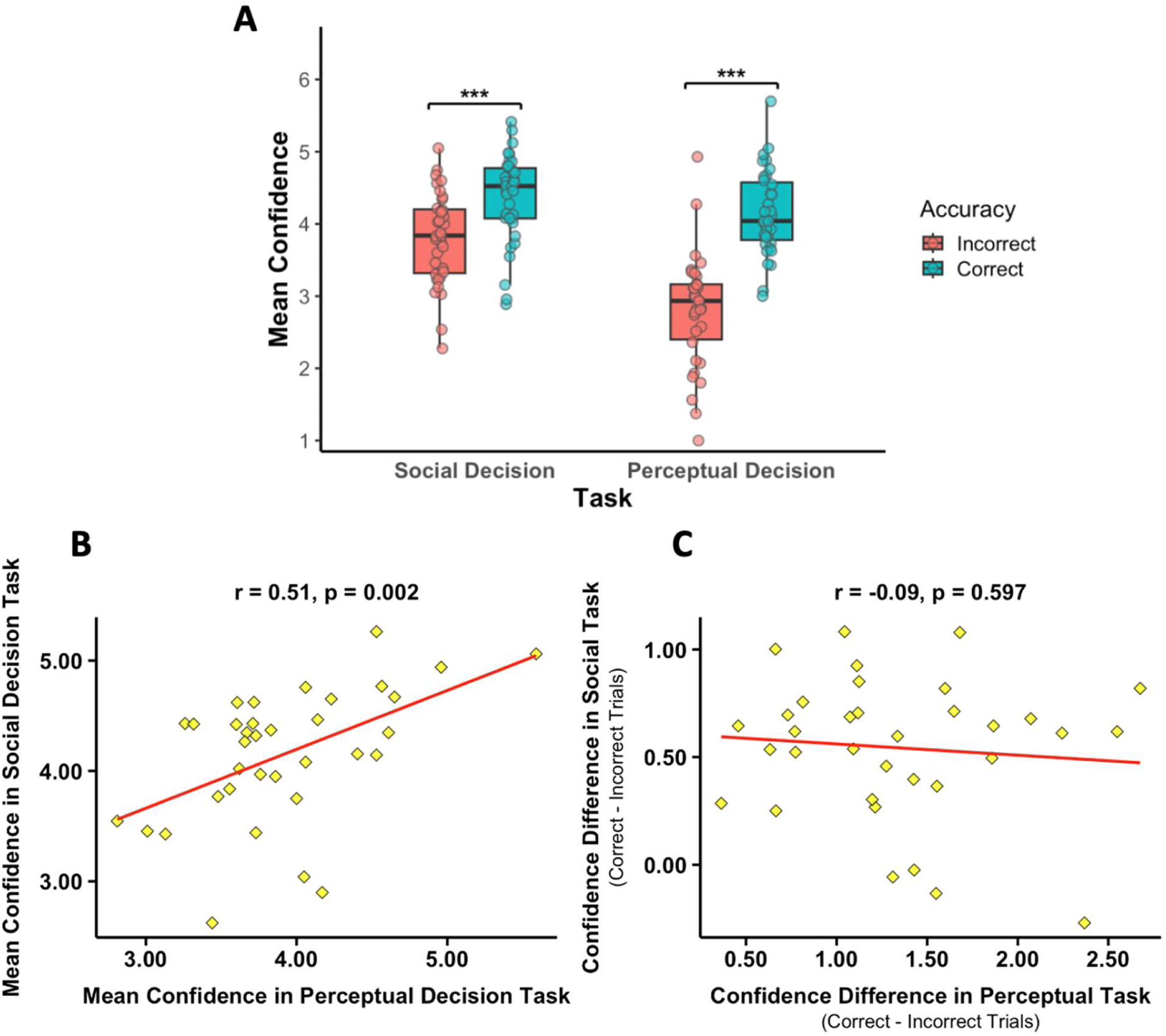
Metacognitive bias and sensitivity in both tasks. (A) Boxplots of confidence ratings for correct and incorrect responses in the social decision task (left) and the perceptual decision task (right) show significant differences in confidence, indicating metacognitive access to the accuracy of participants’ decisions. (B) Scatterplot of metacognitive bias as measured by mean confidence, indicating a significant cross-task correlation of confidence between the social decision task (y-axis) and the perceptual decision task (x-axis). (C) Scatterplot of metacognitive sensitivity, calculated as confidence_correct_ – confidence_incorrect_, in the social decision task (y-axis) and the perceptual decision task (x-axis).

### Overlapping neural networks encode confidence in social and non-social domains

Based on the behavioural findings, according to which participants possessed metacognitive access to the accuracy of their social and perceptual decisions, we compared the neural correlates of metacognitive judgements between these tasks in our fMRI data. We hypothesised that both social and perceptual metacognition are encoded in a domain-general prefrontal core network for metacognition, which may, however, show domain-specific connectivity patterns with areas involved in first-order task processing, in line with two-layer accounts of metacognition. To test these predictions, we first compared the neural correlates of first-level task performance in the social and perceptual decision task: while we found enhanced activation in, among other brain regions, bilateral TPJ, dmPFC, and ventral posterior cingulate cortex (vPCC) during social compared with perceptual decisions, all *p* < 0.01, FWE-corrected at cluster level (Table S1), the brain regions that showed stronger activation during perceptual versus social decisions were the occipital cortex (OC) extending into parietal cortex, ACC, and primary motor cortex. Further significant clusters were the bilateral insula as well as white matter in proximity to the hippocampus, all *p* < 0.03, FWE-corrected at cluster level (Table S2). This suggests that first-order social and perceptual decisions correlated with dissociable brain networks (Fig 3A).

**Fig 3.**
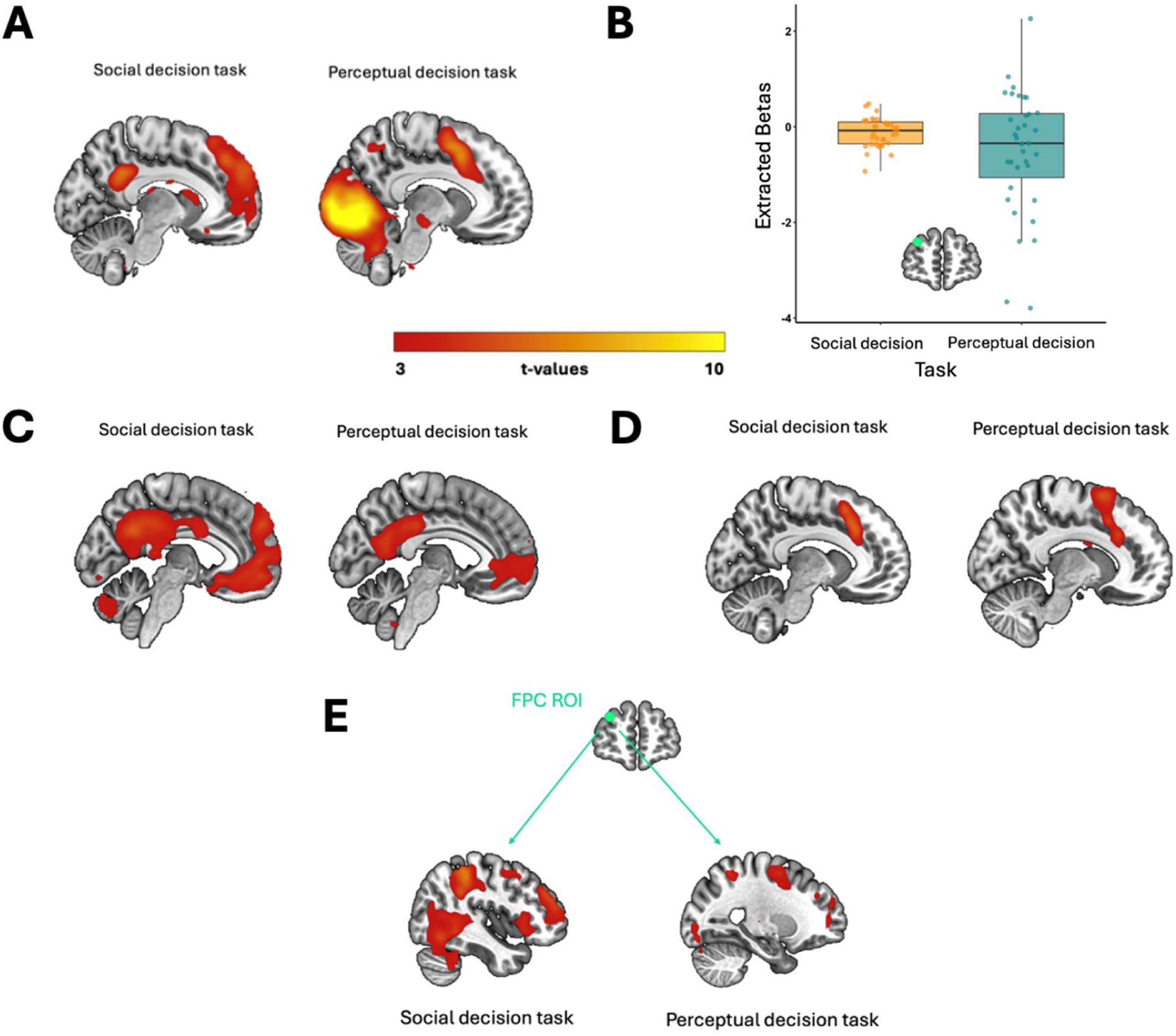
Neural correlates of task performance and confidence in both tasks. (A) First-order task performance showed neural activity in bilateral TPJ and dmPFC in the social decision task (left) and in OC and ACC in the perceptual decision task (right). (B) Extracted parameters obtained with an 8 mm sphere ROI in the FPC for the social decision task (left) and the perceptual decision task (right) reveal significant activation in both tasks during confidence rating. Dots indicate individual data points. (C) Confidence ratings positively correlated with activation in dmPFC in the social decision task (left) and in aPFC in the perceptual decision task (right). (D) Confidence ratings negatively correlated with activation in dmPFC in the social decision task (left) and in the perceptual decision task (right). (E) Connectivity of the FPC as seed region with other brain regions showed task-specific significant activity in the precuneus extending into bilateral TPJ in the social decision task (left) and in OC in the perceptual decision task (right), albeit the latter effect is only significant in the ROI analysis. All activation maps are thresholded at *p* < 0.001, uncorrected.

Next, we assessed whether domain-general or-specific brain regions encode second-order confidence judgements in these tasks. In line with our hypothesis that the FPC represents a domain-general neural substrate of metacognition, we found that the FPC (extracted from an ROI from a meta-analysis (5)) correlated with decreasing confidence judgements both in the social (beta = –0.12, *t*(34) = 2.26, *p* = 0.03) and the perceptual decision task (beta = –0.52, *t*(34) = 2.38, *p* = 0.02; Fig 3B). This FPC activation was robust to controlling for task difficulty, again in both the social (beta = –0.18, *t*(34) = 2.84, *p* < 0.01) and the perceptual decision task (beta = –0.23, *t*(34) = 2.71, *p* = 0.01) (Fig S2). The extracted parameters for confidence-related FPC activation were, however, not correlated between tasks, *r* = 0.14, *p* = 0.43. Exploratory whole-brain analyses revealed that, in the social decision task, confidence was positively correlated with activations in the dmPFC as well as precuneus (Fig 3C, Table S3) and negatively correlated with activations in right anterior dlPFC/dmPFC (Fig 3D, Table S4). These (prefrontal) findings were also robust when controlling for task difficulty. In the perceptual decision task, confidence was positively correlated with activations in the left and right putamen, left FPC, vmPFC, and right ventral posterior cingulate cortex (Fig 3C, Table S5); in contrast, we observed negative correlations between confidence and activations in right dmPFC, ACC, and right insula (Fig 3D, Table S6). A direct comparison between the two tasks revealed that the dmPFC showed significantly stronger representations of confidence in the social compared with the perceptual decision task, whereas confidence representations were stronger in the perceptual relative to the social decision task in bilateral putamen, and right angular gyrus extending into occipital cortex (Tables S7-8). An overlap analysis showed that activations in the angular gyrus and orbitofrontal cortex were positively correlated with confidence in both tasks (Table S9), whereas confidence was negatively correlated with activations in dmPFC and ACC (Table S10). Overall, these findings were in line with previous studies reporting neural activation in the FPC/anterior PFC to encode subjective confidence across domains (6,11,37,38).

### Domain-general and –specific connectivity with FPC during social and perceptual metacognitive judgements

While we observed overlapping networks, including the FPC and dmPFC, to encode confidence in social and perceptual decision making, two-layer accounts of metacognition assume that metacognitive judgements require the FPC to access information about task difficulty represented in task-specific brain areas. Therefore, we performed psycho-physiological interaction (PPI) analyses with the FPC as seed region to examine whether connectivity patterns of the FPC with other brain regions differed between the two tasks. In the social decision task, we found significant coupling of the FPC with bilateral dlPFC, right dmPFC, and bilateral insula for decreasing confidence, all *p* < 0.01 FWE-corrected at cluster level (Table S11). In the perceptual decision task, FPC showed confidence-related coupling with the right dlPFC, dmPFC, inferior frontal cortex, as well as the left posterior parietal lobe, all *p* < 0.05 FWE-corrected at cluster level (Table S12). Again, we performed a conjunction analysis that revealed overlapping confidence-related coupling between FPC and dmPFC (Table S13). However, other functional connectivity patterns with FPC appeared to be domain-specific: In the social task, we found significant confidence-related connectivity of the FPC with a parietal network consisting of precuneus and extending into bilateral TPJ and temporal lobe on the whole-brain level, all *p* < 0.01 FWE-corrected at cluster level (Fig 3E, Table S9), but not in the perceptual decision task. Again, this connectivity between FPC and TPJ/temporal lobe was robust to controlling for task difficulty, all *p* < 0.01 FWE-corrected at cluster level. Note, however, that no areas showed significant differences in confidence-related functional coupling with FPC in a direct comparison between the social and the perceptual decision task, all *p* > 0.05 FWE-corrected at cluster level. While we found no significant connectivity results selectively in the perceptual decision task on the whole-brain level (Fig 3E), an explorative analysis with an ROI (based on the coordinates for the visual cortex (x = –12, y = –85, z = – 8) reported in a previous study on perceptual metacognition (6)) revealed significant connectivity between FPC and visual cortex in the perceptual decision task, ß = –0.55, *t*(34) = 2.43, *p* = 0.02, but not in the social decision task, ß = –0.10, *t*(34) = 1.50, *p* = 0.14 (though we note that the comparison between the tasks showed only a non-significant trend-level effect, ß = –0.44, *t*(34) = 1.81, *p* = 0.08). Thus, in addition to domain-general coupling between FPC and prefrontal areas during metacognitive judgements, there was preliminary evidence for domain-specific coupling with areas involved in first-order task processing, though we note that these imaging results should be interpreted with caution given that the direct comparisons between the social and perceptual domains were not significant.

### Brain stimulation reveals domain-specific networks causally underlying metacognitive sensitivity

While the imaging results suggest both domain-general and domain-specific connectivity patterns for metacognition in social and perceptual decisions, they provide no conclusive evidence for domain-specific neural networks due to a lack of significant differences between the domains; moreover, they do not allow concluding that these connectivity patterns are causally relevant for metacognition due to the correlative nature of neuroimaging findings. We therefore conducted two further experiments to address these issues: In particular, we tested whether synchronising (in-phase tACS) versus de-synchronising (anti-phase tACS) theta activation between FPC and the TPJ (Experiment 2) and the OC (Experiment 3) leads to domain-specific modulations of social and perceptual metacognition, respectively. We chose the TPJ and OC as domain-specific areas that might be coupled to the FPC core network during metacognitive judgements based on our PPI results in the imaging experiment. Participants performed the same experimental tasks as in the imaging experiment while receiving either in-phase, anti-phase, or sham tACS in separate blocks (within-subject design). Based on the assumption of domain-specific neural networks for metacognition, we hypothesized bifocal FPC-TPJ and FPC-OC tACS to show domain-specific effects on metacognitive sensitivity during social and perceptual decisions.

In both tACS experiments, we observed no significant effects of in-phase or anti-phase (versus sham) tACS on first-order task performance in the social decision task, all *p* > 0.08, or the perceptual decision task, all *p* > 0.46. We did, however, find evidence for domain-specific modulations of metacognitive sensitivity (expressed by increased confidence after correct compared with incorrect decisions): In the FPC-TPJ experiment, a GLMM regressing confidence on predictors for tACS and Accuracy revealed that metacognitive sensitivity in the social decision task was marginally increased under in-phase compared with sham tACS (In-phase × Accuracy, ß = 0.10, *t*(39)= 1.99, *p* = 0.05), but not under anti-phase compared with sham tACS (Anti-phase × Accuracy, ß = –0.004, *t*(96) = 0.10, *p* = 0.92) (Fig 4A). A direct comparison between the active stimulation conditions revealed significantly better metacognitive sensitivity under in-phase versus anti-phase tACS, indicating a phase-specific effect (Anti-phase × Accuracy, ß = –0.11, *t*(49) = 2.28, *p* = 0.03). In the perceptual decision task, in contrast, neither in-phase (In-phase × Accuracy, ß = –0.01, *t*(35) = 0.23, *p* = 0.82) nor anti-phase tACS (Anti-phase × Accuracy, ß = –0.02, *t*(35) = 0.32, *p* = 0.75) led to significant changes in metacognitive sensitivity (Fig 4B). This suggests that strengthening the FPC’s coupling with the TPJ changed the accuracy of metacognitive judgements specifically in the social domain. The reversed result pattern occurred under FPC-OC stimulation: While we found no significant tACS effects on metacognitive sensitivity in the social decision task (In-phase × Accuracy, ß = 0.02, *t*(107) = 0.27, *p* = 0.79; Anti-phase × Accuracy, ß = 0.009, *t*(114) = 0.23, *p* = 0.82) (Fig 4C), we observed a significant reduction of perceptual metacognitive sensitivity for anti-phase versus sham tACS (Anti-phase × Accuracy, ß = –0.10, *t*(86) = 2.24, *p* = 0.03) (Fig 4D). In-phase versus sham tACS showed no effects on perceptual metacognition (In-phase × Accuracy, ß = –0.07, *t*(42) = 1.40, *p* = 0.17). A direct comparison between the active stimulation conditions did not reveal a significant difference in metacognitive sensitivity (Anti-phase × Accuracy, ß = –0.04, *t*(144) = 0.90, *p* = 0.37). The lack of significant main effects of tACS in these analyses, all *p* > 0.14, suggests that the stimulation affected metacognitive sensitivity specifically, rather than confidence judgements per se (metacognitive bias). To more thoroughly assess whether FPC-TPJ versus FPC-OC tACS modulated metacognitive sensitivity in a domain-specific way, we ran another GLMM comparing in-phase and anti-phase versus sham tACS, which combined both experiments and both tasks. This analysis suggested marginally significant dissociable effects of anti-phase versus sham tACS on metacognitive sensitivity in the social versus perceptual domain (Experiment × Task × Accuracy × Anti-phase tACS: ß = –0.34, *t*(79) = 1.96, *p* = 0.05). Importantly, a direct comparison between in-phase versus anti-phase tACS revealed a significant four-way interaction (Experiment × Task × Accuracy × Anti-phase tACS: ß = –0.43, *t*(72) = 2.24, *p* = 0.03). Together, this suggests that domain-specific functional connectivity patterns may causally implement metacognitive sensitivity in the social (FPC-TPJ connectivity) and perceptual (FPC-OC connectivity) domains.

**Fig 4.**
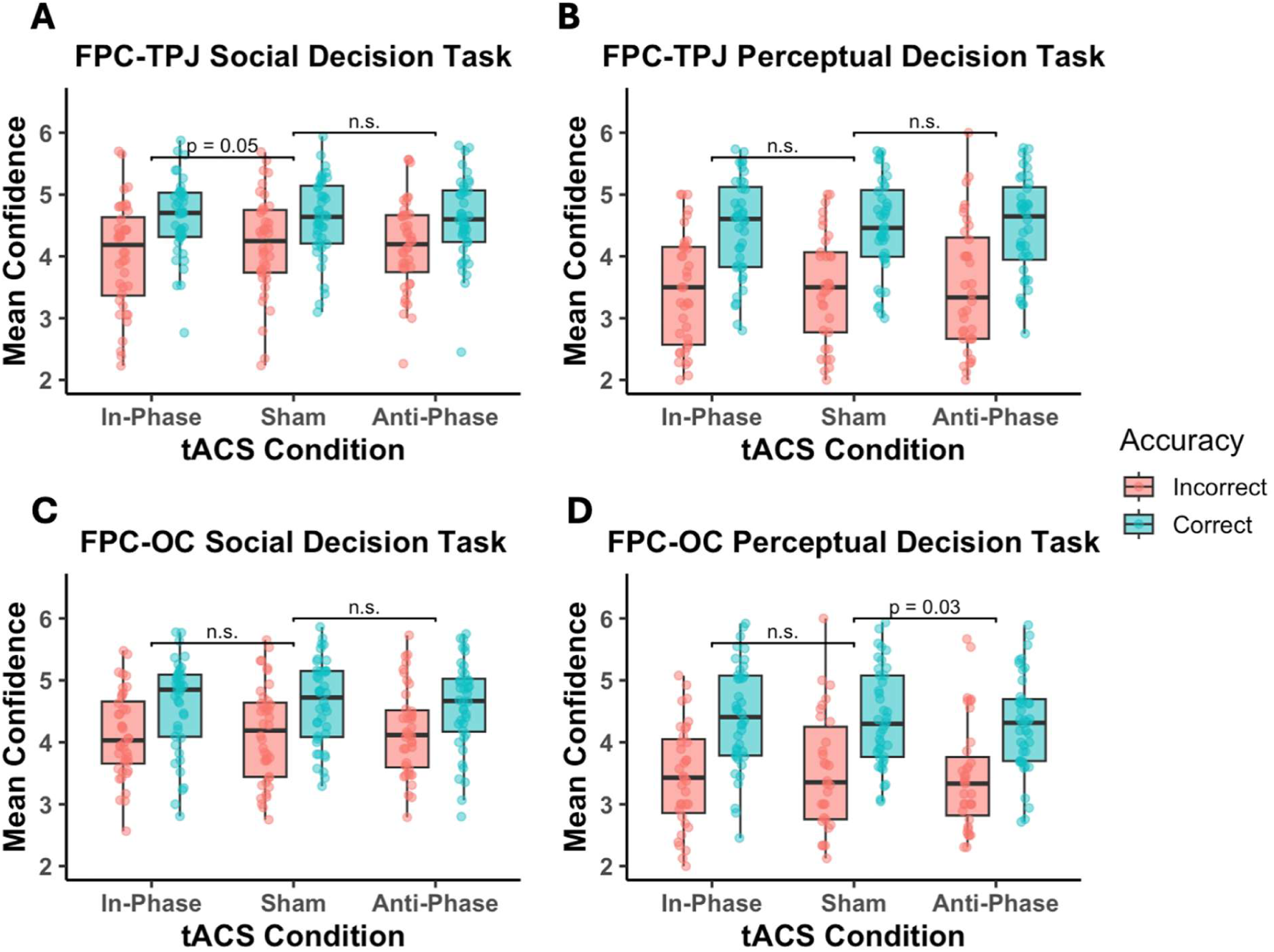
Stimulation effects on metacognitive sensitivity. Stimulation effects on mean confidence separately for correct versus incorrect trials and for all three tACS conditions (in-phase, sham, anti-phase). In Experiment 2, FPC-TPJ in-phase tACS marginally modulated metacognitive sensitivity compared with sham and significantly compared with anti-phase in the social decision task (A) but not in the perceptual decision task (B). In Experiment 3, FPC-OC tACS did not have an effect in the social decision task (C), but anti-phase versus sham tACS significantly decreased metacognitive sensitivity in the perceptual decision task (D).

Based on the finding that manipulating the FPC’s coupling with brain areas for first-order task processing changes metacognitive sensitivity (while leaving first-order task performance or metacognitive bias unaltered), we next aimed to elucidate the computational mechanisms underlying this effect. Improved metacognitive sensitivity might be associated with higher metacognitive efficiency, or result from lower metacognitive noise, impeding the computation of confidence signals. While metacognitive efficiency puts a person’s metacognitive skills in relation to her first-order task performance (Mratio), metacognitive noise refers to the distortion of confidence read-outs through internal computational processes. When we analysed participants’ individual Mratio scores (computed separately for task and tACS condition), this did not reveal significant stimulation effects on metacognitive efficiency, all *ps* > 0.08. However, when testing for stimulation effects on metacognitive noise, we found that anti-phase versus sham FPC-TPJ tACS increased metacognitive noise in the social decision task (ß = 0.48, *z* = 2.10, *p* = 0.04), whereas in-phase versus sham stimulation did not show a significant effect (ß = –0.03, *z* = 0.12, *p* = 0.91) (Fig 5A). A direct comparison between in-phase versus anti-phase tACS revealed a marginally significant tACS effect on metacognitive noise, ß = –0.5, *z* = 1.91, *p* = 0.06. In the perceptual decision task, we did not find significant FPC-TPJ tACS effects (In-phase versus sham tACS: ß = 0.09, *z* = 0.43, *p* = 0.67; Anti-phase versus sham tACS: ß = 0.20, *z* = 0.97, *p* = 0.33) (Fig 5B). Thus, in analogy to the findings for metacognitive sensitivity, FPC-TPJ stimulation reduced metacognitive noise selectively in the social domain. For FPC-OC stimulation, however, we found no stimulation effect, neither in the social task (Fig 5C) nor in the perceptual task (Fig 5D), all *p* > 0.10, in contrast to the findings for metacognitive sensitivity with this stimulation setup. Nevertheless, when we assessed whether stimulation effects on metacognitive noise differ between FPC-TPJ and FPC-OC tACS, we found a significant In-phase versus Anti-phase tACS × Task × Experiment interaction, ß = –1.12, *z* = 2.13, *p* = 0.03, suggesting that FPC-TPJ versus FPC-OC tACS yielded domain-specific effects on metacognitive noise. Overall, these results show that changing the FPC’s synchronous activation with regions involved in first-order task processing influences metacognitive sensitivity and noise in confidence computations in a domain-specific way.

**Fig 5.**
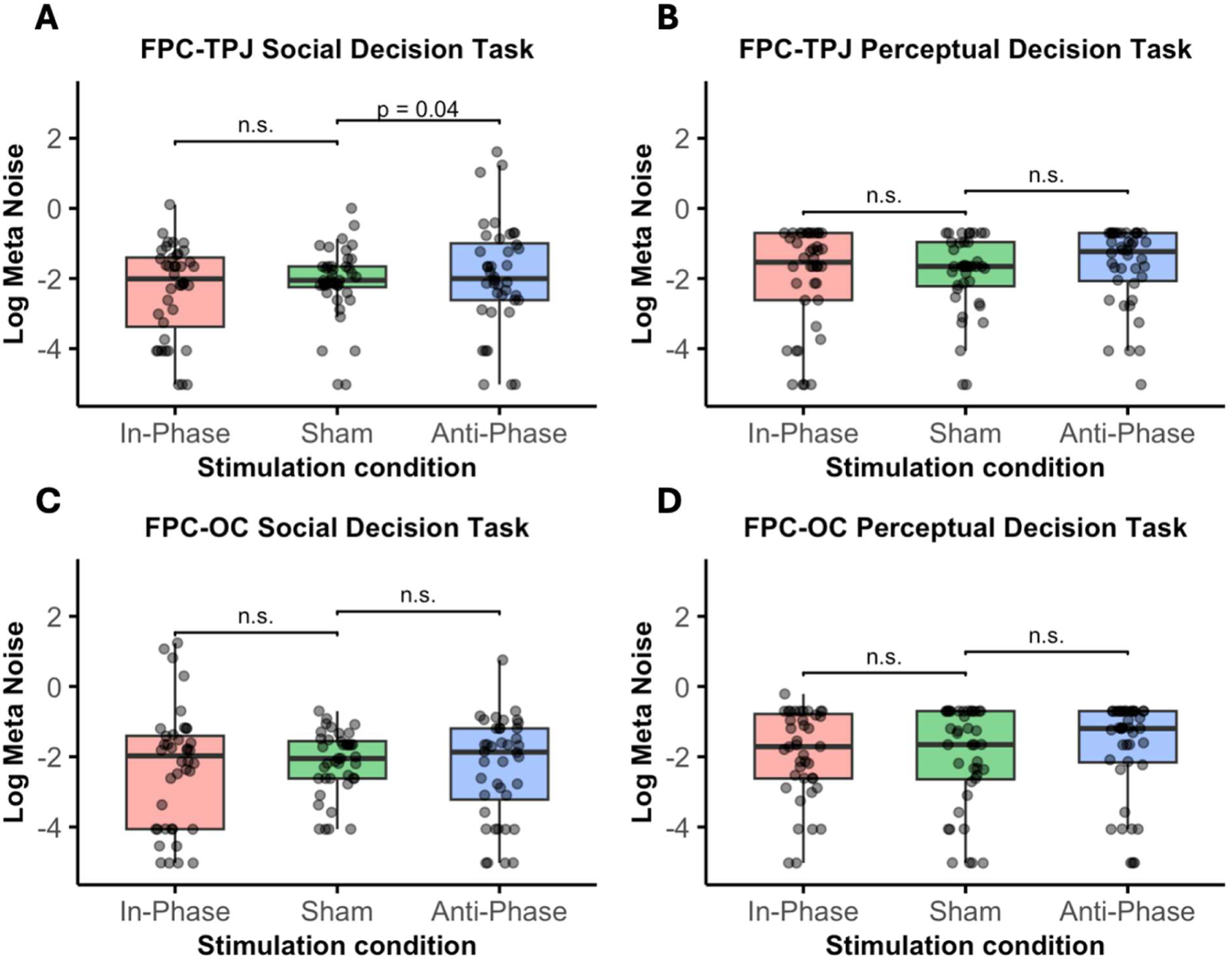
Stimulation effects on metacognitive noise. Stimulation effects on log-transformed metacognitive noise for all three tACS conditions (in-phase, sham, anti-phase). In Experiment 2, FPC-TPJ tACS showed increased metacognitive noise under anti-phase compared to sham tACS in the social decision task (A), but not the perceptual decision task (B). In Experiment 3, FPC-OC tACS did not show stimulation effects on metacognitive noise in neither the social decision task (C) nor the perceptual decision task (D).

## Discussion

The question of whether metacognition is domain-general or domain-specific has not yet been conclusively answered. Therefore, the present study aimed to shed light on this question from a neuroscientific perspective by examining whether common or dissociable neural mechanisms underlie metacognition in social and perceptual domains. Based on two-layer accounts, we predicted a prefrontal core network for metacognition to be functionally coupled with brain areas for mentalising in a novel task measuring social metacognition, with the brain networks underlying social metacognition differing from those underlying perceptual metacognition. On the behavioural level, we observed moderate correlations between metacognitive processes in social and perceptual decisions, consistent with previous findings for other domains (20). Similarly to recent empirical evidence (39), we found these correlations to be moderate in confidence ratings, but not present in metacognitive sensitivity, efficiency, or noise. We note that tendencies to report low confidence were speculated to reflect personality traits (40), which is consistent with the observed correlation across domains, though confidence ratings are also influenced by contextual factors like mood (41). The distinction between general and specific components was mirrored by the neural findings: On the one hand, several brain regions, particularly in the prefrontal cortex, seem to be involved in both social and perceptual metacognition, such as FPC, dmPFC, and dlPFC. On the other hand, we found domain-specific connectivity of FPC with areas involved in first-order social and perceptual processing (TPJ and OC, respectively). Overall, our findings provide a nuanced perspective on the generality versus specificity debate of metacognition, and support neural two-layer accounts according to which metacognition relies on the communication between domain-general prefrontal areas and domain-specific areas involved in task performance.

For both metacognitive tasks, we found functional overlap in prefrontal areas, such as FPC and dmPFC. This aligns with the existing neuroimaging literature on metacognition, where FPC has been associated with metacognitive judgements across a range of tasks (5,7,10). It belongs to a larger frontoparietal network that has been ascribed the role of an integratory hub of different inputs, being at the top of the hierarchy of prefrontal control processes (42,43). To make confidence judgements about performance in a task, the FPC requires access to information about task difficulty, which might be represented in both domain-general (dmPFC, dlPFC), and domain-specific brain regions (TPJ, OC). Therefore, given the behavioural correlations we found between social and perceptual metacognition, FPC’s involvement in several sub-domains concurs with the (hypothesised) integrative function of this core network. Additional brain regions that appear to overlap in their functional involvement in metacognition are dmPFC and dlPFC, which are associated with metacognitive performance: Evidence from brain stimulation studies suggests that dmPFC is causally involved in metacognitive bias (8) and that dlPFC reads out sensory signal strength to relay it to hierarchically higher brain regions such as anterior PFC (9). In the past, these brain regions have been mentioned as forming parts of the decision network (10,44). In both metacognitive domains, we moreover found significant connectivity between FPC and dmPFC, as well as dlPFC. It has been suggested that accurate metacognitive judgements rely on cross-talk between FPC and the decision network, such that the former may read out and integrate information from the latter. In a recent study that combined brain stimulation and neuroimaging (11), the specific connectivity pattern between FPC and dlPFC has been found to causally guide metacognition, thereby confirming the important role of cross-talk between FPC and the decision network (dlPFC and dmPFC).

In addition to the shared confidence representations in the prefrontal network, we found domain-specific connectivity patterns of FPC with other brain regions involved in first-order task processing. For the social metacognition task, we found connectivity between FPC and TPJ, which has long been established as a functional hub for social cognition, such as mentalising (36). In the perceptual task, FPC showed connectivity with the visual cortex, consistent with previous findings (6), though note that this effect was present only in an ROI analysis and did not significantly differ between the domains. While the domain-specificity of the PPI results therefore needs to be interpreted with caution, this correlational evidence was further strengthened by results from our two tACS experiments, which showed that bifocal tACS causally influenced metacognitive sensitivity in a dissociable, domain-specific manner. Modulating connectivity between task-relevant brain regions (TPJ vs. OC) and FPC not only affected metacognitive sensitivity in the given task, but also showed causal influence on metacognitive noise, an internal variability contributing to metacognitive inefficiency (45). Note that while we found this effect on metacognitive noise only for the social decision task in Experiment 2, it provides first empirical evidence for a neural basis of metacognitive noise, suggesting that noise in confidence read-outs may depend on FPC’s connectivity with areas involved in first-order task processing. The stimulation-induced changes in metacognitive noise may have resulted in the observed tACS effects on metacognitive sensitivity, given that noisier confidence representations weaken the link between performance accuracy and metacognitive confidence reports (46).

These differential findings support the theoretical two-layer architecture of metacognition (29,30), wherein the prefrontal regions, such as the FPC, read out decision-relevant information provided by task-related brain regions. The regions we find here are differentially involved in the first-order tasks, respectively. Therefore, these findings suggest that while neural evidence can be found for the behavioural correlation in the form of overlapping brain regions in prefrontal areas, there are also differential connectivity patterns that pertain to each sub-domain in a domain-specific way. These insights extend recent neuromodulation findings showing that stimulating sensory brain areas impairs both first-order and second-order (metacognitive) performance, whereas stimulating more meta-level brain regions affects metacognition alone, leaving first-order task performance intact (47). This co-existence of both domain-general and domain-specific elements in the underlying neural architecture is in line with a recent meta-analysis on metacognition across domains (27).

Nevertheless, some of our findings should be interpreted with caution, owing to the following limitations: First, the lack of a tACS effect on metacognitive noise in Experiment 3 introduces some inconsistency into the pattern of the rest of our results. It is, however, in alignment with the somewhat weaker PPI results in the perceptual task compared to the social task we found in the neuroimaging data. Since it is a broad network of occipital and parietal areas that is functionally involved in visual perception and has previously been shown to be coupled with the FPC during metacognitive judgements (6), there might be other areas of the network that are more strongly functionally coupled with the FPC. Alternatively, the occipital areas involved in perceptual metacognition processing might lie at a more subcortical region than the ones in the FPC, which could explain why the specific tACS setup in the current study did not sufficiently modulate activation in the targeted region. Nevertheless, the significant difference between tasks and experiments we report clarifies the domain-specificity of the FPC-TPJ stimulation effect on metacognitive noise.

Second, given the tACS effects on metacognitive sensitivity and noise, the lack of a tACS effect on Mratio as a measure of metacognitive efficiency in either experiment may appear surprising. Mratio quantifies metacognitive skills relative to individual first-order task performance, such that subtle and non-significant stimulation effects on social and perceptual decisions may have resulted in the lacking tACS effects on metacognitive efficiency (keeping in mind that in our bifocal tACS setup the TPJ and OC as areas for first-order task performance were co-stimulated in Experiment 2 and 3, respectively). Alternatively, significant effects on metacognitive sensitivity but not efficiency were also observed in other studies (3,48) and were interpreted as evidence for a “mapping problem” (i.e., performance error detection is intact but not translated into aligned confidence reports). In contrast, changes in metacognitive efficiency would reflect a “detection problem” where individuals have difficulties in metacognitively detecting performance errors. In any case, the observed discrepancy between metacognitive sensitivity/noise and efficiency falls within the line of argument brought forth by Shekhar & Rahnev (46) for the dependence of Mratio on participants’ metacognitive bias. This observation led them to propose a new model of metacognition including metacognitive noise that should be independent of one’s metacognitive bias. Following this, the lack of tACS effects on Mratio might be attributed to a possible confound between this measure and metacognitive bias. In any case, the current findings suggest that phase-specific coupling between FPC and areas related to first-order task processing influences metacognitive noise, which may result in changes in metacognitive sensitivity.

Our results have implications for clinical populations involving impaired metacognition. There is recent evidence for a domain-general account of metacognition in depression and obsessive-compulsive disorder (49,50). However, exploring social metacognition would be especially interesting for clinical populations characterised by difficulties with social cognition, such as autism spectrum disorder or borderline personality disorder. For these clinical populations, elucidating whether certain domain-specific deficits on the behavioural level might be related to suboptimal domain-specific metacognitive processes could be especially important, since ultimately, this could pave the ground for the development of specifically targeted interventions to improve metacognition or prevent it from deteriorating.

## Conclusion

This study shows moderate correlations between social and perceptual metacognition that are mirrored in their underlying neural correlates: we find that common brain regions, such as the FPC, dmPFC, and dlPFC, are functionally involved in both metacognitive tasks.

Alongside this, we find that connectivity between the FPC and brain regions involved in first-level task performance (TPJ and OC, respectively) causally underlies metacognition in a domain-specific way. Thus, our results serve as evidence for two-layer accounts, suggesting that metacognition relies on a co-existence of both domain-general and domain-specific signals in the human brain.

## Materials and Methods

### Experiment 1 (fMRI) Participants

The imaging experiment’s sample included 35 participants (14 female, 33 right-handed, mean age = 24.43 years, range 18-34 years), following an a priori power analysis requiring 35 participants for a moderate effect size of Cohen’s d = 0.50 (power = 80%, alpha = 5%) for the correlation between confidence and prefrontal activation. Before the experiment, participants confirmed they had normal or corrected vision, no psychiatric and/or medication history, no metallic implants, and gave written informed consent. Participation was compensated with 20€/h and an additional bonus based on a randomly picked trial outcome. An additional 20 volunteers (11 female, mean age = 25.25, range 21-32 years) participated in a behavioural pilot experiment (outside the MRI scanner) and were compensated with 15€/h. The study was conducted in accordance with the ethical standards laid down in the Declaration of Helsinki and approved by the ethics committee of the department for psychology at the Ludwig-Maximilians-Universität München (protocol number: “20_Soutschek_a”).

### Stimuli and Task Design

Participants performed two tasks inside the MRI scanner: a perceptual decision task and a social decision task. Both of these tasks were programmed in MATLAB, R2023) using Psychtoolbox (51).

#### Perceptual Decision Task

The perceptual decision task followed the paradigm developed by Fleming et al. (52) and the variant used in a previous study in our lab (8). In each trial, participants were shown two white circles (diameter 5.1°), each containing three crosses, together with a fixation cross on a screen with a black background (0.5 s), followed by the same circles containing a variable, unequal number of dots (diameter 0.1°) for 0.7 s.

Participants were asked to indicate which of the circles contained more dots within 2 s by pressing the left or right button for the corresponding response. The number of dots inside each circle was bounded to 1–100, and the contrast was adjusted to participants’ individual performance level, which was determined in a previous practice task outside the MRI scanner using a one-up two-down staircase procedure (52,53). The staircase procedure aimed to balance the difficulty of the task between individuals, thus controlling for individual performance differences. After each choice and a jittered inter-stimulus interval (mean duration = 1 s), participants had to report their subjective confidence that their answer was correct on a rating scale ranging from “low” to “high” in six steps and randomised per trial in scale orientation (low-high vs. high-low) within 3 s (Fig 1A). The next trial started after a jittered inter-trial interval of 1 s.

#### Social Decision Task

The social decision task was based on a variant of the Ultimatum Game (54). Participants were told that they were playing in the role of the proposer and would see pre-defined offers on how to split an amount of twenty coins between themselves and anonymous responders (e.g., You: 8 coins and Other: 12 coins). To make the social and the perceptual decision tasks as visually similar as possible, the offers were shown in circles that matched the perceptual decision task (see below). Following the rules of the Ultimatum Game, the responder had to choose between accepting and rejecting the offer. Participants were made aware that both they, as proposers, and the responders would only receive the coins if the responder accepted the offer. We informed participants that the responders’ decisions had been made by real participants, as they stemmed from a previous study in our lab (55). Participants’ task was to predict whether the anonymous responders would accept or reject a given offer in each trial. The choice options “accept” and “reject” were presented below the offers on the left and right screen side (in counterbalanced order) and participants indicated their choice by pressing a left or right button, respectively, on an MRI-compatible button box within 4 s. After each choice, the selected option turned red and was shown together with the offer for the remaining time of the 4 s. After a Poisson-distributed jittered inter-stimulus interval (mean duration = 1 s), participants were presented with a sliding scale to indicate their confidence in having given the correct prediction (Fig 1B). As for the perceptual decision task, the scale ranged from 1 (low confidence) to 6 (high confidence), and participants had to select their level of subjective confidence within 3 s. The orientation of the confidence scale was randomised. Then, participants received feedback about whether the responder had accepted or rejected the offer (duration = 1 s) based on the decisions of real responders in our previous study. We chose the real behavioural data from different participants in our previous study (55) at random to minimize learning effects in this task. The next trial started after a Poisson-distributed jittered inter-trial interval with a mean duration of 4 s.

### Procedure

Participants carried out several practice trials (50 for the perceptual and 10 for the social decision task) outside the scanner to familiarise themselves with the tasks. Specifically for the perceptual decision task, the practice trials served as a staircase procedure to determine the individual’s first-level performance. Based on this, the task difficulty inside the scanner in the main experiment was adjusted to each participant’s individual performance. Next to the practice trials, participants filled out several questionnaires, the Autism Questionnaire (AQ, 56), the UCLA Loneliness Scale (57), and they also performed the Culture-Fair test (CFT 20-R, 58) IQ test. These measures will be reported in a separate manuscript on metacognition in autism. Inside the scanner, they performed 5 runs, with each run including 20 trials for the perceptual decision and 20 trials for the social decision task presented in a randomised order at the trial level. Each run lasted approximately 8 min and was followed by a short break inside the scanner. Participants performed the first three runs before the T1-scan for the structural images, during which they had time to take a break and relax. In total, participants spent approximately 50 min inside the MRI scanner.

### MRI data acquisition

The functional MRI data were obtained using a 3 T Siemens PRISMA MRI scanner with a 64-channel head coil. Functional scans were acquired with a T2*-weighted gradient-echo, echo-planar pulse sequence in ascending interleaved order (3 mm slice thickness, 3.0 × 3.0 mm in-plane resolution, 70 × 70 voxels per slice, flip angle = 45◦, FOV = 210). Echo time (TE) was set to 30 ms, and repetition time (TR) was 1000 ms. A total of 48 axial slices were obtained (slice gap = 10%). A T1-weighted 3D MPRAGE image was acquired for anatomical reference (0.8 mm slice thickness, voxel size = 0.8 × 0.8 × 0.8 mm, 208 sagittal slices, flip angle = 8◦, TE = 2.22 ms, TR = 2500 ms).

### Behavioural data analysis

We analysed behavioural data with general(ised) linear mixed models (GLMMs) in R (59) using the packages lme4 (60) and lmerTest (61). In all analyses, the alpha threshold was set to 5% (two-tailed). Trials where participants had not reacted were excluded from the analysis. Across all experiments, this affected a mean percentage of 1.57% of all trials. In all experiments, there was sufficient variation in participants’ confidence ratings (i.e., all participants reported at least four unique values on the confidence scale) in both tasks.

We compared performance accuracy between the two tasks with a GLMM regressing accuracy on a categorical fixed-effects predictor for Task (0 = perceptual task, 1 = social task). Similarly, in order to analyse metacognitive bias, we ran a GLMM regressing confidence on a categorical fixed-effects predictor for Task (0 = perceptual task, 1 = social task). For metacognitive sensitivity, we ran separate GLMMs regressing confidence on a categorical fixed-effects predictor for Accuracy (0 = incorrect, 1 = correct). As an additional measure, we calculated mean confidence differences between correct and incorrect trials ((mean confidence)_correct_ – (mean confidence)_incorrect_). For the analysis of metacognitive efficiency, we computed individual Mratio values using the *metaSDT*-package (62) with the SSE method. As a further measure of metacognition, we computed individuals’ metacognitive noise, which models the internal corruption added to confidence judgements, isolating metacognitive inefficiency from task performance and confidence bias (46). We computed metacognitive noise following the computational approach reported by Rahnev (63). See the Supplementary Material (Note S1) for more information on the model. For all these measures, we calculated between-task correlations with the Pearson method.

### Imaging analysis

The fMRI analyses were performed using SPM12 (Version 7487, Wellcome Department of Cognitive Neurology, London, UK) implemented in MATLAB R2023 (MathWorks Inc., Natick, MA, USA).

For the preprocessing, each participant’s functional images were realigned and unwarped to correct for head motion and susceptibility-by-movement effects. Slice-timing correction was then applied using the ascending interleaved acquisition order. The T1-weighted anatomical image was coregistered to the mean functional image and segmented into tissue classes. The data were subsequently normalised to MNI space. Spatial smoothing was carried out with an 8 mm FWHM Gaussian kernel, and for first-level model specification, temporal filtering was applied using a high-pass cutoff period of 128 seconds.

For the statistical analysis, we computed two first-level general linear models (GLMs) for each participant. Both GLMs included 4 task-related onset regressors: for the social decision task, we modelled one onset regressor for the social decision screen (first-order task performance) with a duration of 4 seconds and a second onset regressor for the post-decision confidence rating (duration = 4 seconds). Similarly, in the perceptual decision task, the duration of the onset regressor for the perceptual decision (first-level task performance) was set to 1 second, and the duration of the onset regressor for the confidence rating phase was set to 4 seconds. In the first GLM (GLM1), we added parametric modulators for trial-level confidence ratings to the onset regressors for the decision phase in the social and perceptual decision task. The second model (GLM2) included an additional parametric modulator (without orthogonalization) to disentangle neural representations of confidence from task difficulty. A total of 6 additional predictors pertained to movement regressors. Onsets of the four regressors were set to their respective screen appearance. From these first-level GLMs, we computed individual contrasts for confidence ratings in the social and perceptual decision task, as well as for the comparison between first-order task performance in these tasks. Participants’ individual contrasts were then entered into second-level random-effect analyses (one-sample t-tests) to test for significant effects on the group level. To control for alpha error inflation, we corrected for multiple comparisons with family-wise error (FWE) correction at the cluster level (alpha = 0.05), using a cluster-inducing threshold of *p* < 0.001 uncorrected. In addition, we performed region of interest (ROI) analyses with the marsbar toolbox (64) for the FPC, given its canonical role for metacognition. The spherical FPC ROI (centred at MNI coordinates: x = 28, y = 50, z = 26; diameter = 8 mm) was defined based on the MNI coordinates reported in the meta-analysis by Vaccaro & Fleming (5), following the procedures from a previous imaging study on metacognition in our lab (11).

Further, we conducted psychophysiological interaction (PPI) analyses to determine connectivity patterns during the two tasks. For this, we extracted the time course of the BOLD signal from the FPC ROI as a seed region and convolved it with confidence ratings in the social and perceptual decision tasks (psychological terms) to calculate task-specific PPI terms. The BOLD signal time course, the psychological terms, and the PPI interaction terms were then added to GLM1 and GLM2, respectively. We tested for significant activation related to the PPI terms with one-sample t-tests on the second level, again using FWE correction at the cluster level. Given previous findings that FPC is coupled with the visual cortex during perceptual metacognition, we defined a spherical ROI (diameter = 8 mm) for the occipital cortex centred at the peak coordinates (x = –12, y = –85, z = –8) observed in a previous study (6).

### Experiment 2 and 3 (tACS) Participants

For the tACS experiments, two groups of forty healthy participants (Experiment 2: 27 female, 35 right-handed, mean age = 24.68 years, range = 21 – 34; Experiment 3: 29 female, 38 right-handed, mean age = 24.75 years, range = 19 – 32) were recruited following a power analysis according to which a minimum of N = 29 is required for each experiment to reach 80% power with an effect size of Cohen’s d = 0.54 from a previous tACS study on metacognition (Soutschek et al., 2021). Participants underwent either FPC-TPJ (Experiment 2) or FPC-occipital cortex (OC) stimulation (Experiment 3). All participants reported normal or corrected-to-normal vision as well as no history of psychiatric or neurological disorders or metal implants. Before taking part, all participants provided written informed consent and received either 15€/h or course credit as compensation. The study was conducted in accordance with the Declaration of Helsinki (65) and was approved by the LMU ethics committee (protocol number “19_Soutschek_a”).

### Stimuli and Task Design

Participants performed the same two tasks, namely the perceptual and social decision task, as in the fMRI experiment (see above for details).

## Procedure

At the beginning of the experiment, participants gave informed written consent before reading the instructions for both tasks and performing 60 practice trials (50 for the perceptual and 10 for the social decision task) to familiarise themselves with the tasks. Then, the tACS setup was prepared. The main experiment consisted of five runs, each with three tACS mini-blocks that were separated by 30-s breaks to avoid after-effects of the previous stimulation (7,66,67). Each mini-block consisted of 20 trials, with each task making up half of the trials and randomised on a trial-by-trial basis. Each of the mini-blocks was either performed under in-phase (synchronisation), anti-phase (desynchronisation), or sham tACS, while the order of the stimulation condition was pseudo-randomised and carried out unbeknownst to the participant. All mini-blocks had a duration of 2.30 min, thereby not surpassing the maximum stimulation duration of 30 min according to the tACS safety guidelines (68).

### tACS protocol

tACS was applied with a high-definition (HD) set-up (69) using a 16-channel stimulator (DC-Stimulator MC, neuroConn, Ilmenau, Germany). In-phase tACS synchronises activity, thereby increasing cortical connectivity between the target regions, whereas anti-phase tACS decreases this connectivity (70). In both the FPC-TPJ condition and the FPC-OC condition, the active electrode over the FPC was placed at AF4 with two reference electrodes at positions AFz and F6 following the international 10-10 system to target the FPC subregion (MNI coordinates, x = 28, y = 50, z = 26) reported in a meta-analysis by Vaccaro & Fleming (5). In the FPC-TPJ condition, we placed the active electrode at CP6 and reference electrodes at C6 and P6 in line with previous TPJ stimulation studies (71,72). In the FPC-OC condition, Oz served as the active electrode, while reference electrodes were placed at PO7 and PO8 to target the occipital pole, following the peak coordinates for the visual cortex (x = –12, y = –85, z = –8) observed in a previous study (6). Circular rubber electrodes with a 2-cm diameter were applied using Ten20 conductive paste (Ten20 EEG Conductive Paste, Weaver and Company) and secured with a head fixation band. In the in-phase condition, tACS was administered to both target regions with an in-phase theta band frequency of 5 Hz, whereas for the anti-phase condition, the target regions were stimulated with anti-phase theta tACS. We used a 5 Hz theta tACS given that communication between distant brain areas is thought to rely on theta oscillations (73). The intensity was set to 1.5 mA peak-to-peak. In the sham condition, the synchronisation condition was applied for the first 15 s, before the stimulation was turned off for the remainder of the mini-block. The ramp-up time of stimulation during each mini-block was set to 6 s, while the ramp-down time was 3 s. After each mini-block participants were asked to rate their perceived discomfort and visual flicker they had experienced in the current mini-block on two scales (1–6). While stimulation did not affect visual flicker ratings, the discomfort ratings across participants were increased during the active stimulation conditions. We therefore include flicker and discomfort ratings as fixed-effects predictors in all following analyses. Before the experiment, we performed electric field simulations using the SimNIBS 4.6.0 (74) with the Ernie example head model. ROI analyses showed setup-specific targeting: the FPC-TPJ setup produced stronger fields in the TPJ than in the OC ROI, whereas the FPC-OC montage produced stronger fields in the OC than in the TPJ ROI, in both phase conditions (Fig S1).

### Behavioural data analysis

We first compared first-level task performance accuracy between stimulation conditions separately for each task by regressing performance accuracy on dummy-coded fixed-effects predictors for tACS (in-phase versus sham, anti-phase versus sham). We modelled tACS also as random slopes in addition to participant-specific random intercepts. As control variables, we included (z-transformed) fixed-effects continuous predictors for self-reported discomfort and perceived visual flicker measured at the end of each mini-block.

For the analysis of metacognitive sensitivity, we ran separate GLMMs for each task and tACS experiment (FPC-TPJ and FPC-OC). For this, we regressed subjective confidence on dummy-coded fixed-effects predictors for tACS, Accuracy (0 = incorrect, 1 = correct), and the interaction term. These predictors were also included as random slopes in addition to participant-specific random intercepts. As control variables, we again included fixed-effects for discomfort and visual flicker ratings as well as for task difficulty. In the perceptual task, difficulty was defined as the contrast value between numbers of dots in each circle for a given trial, i.e., higher contrast values represent easier trials. For task difficulty in the social task, we calculated each individual’s indifference point and, for each trial, the distance from this value. Therefore, higher distance values indicate easier trials. Task difficulty was included as a z-scaled predictor. We also ran these separate GLMMs directly comparing in-phase versus anti-phase tACS, in order to control for non-specific stimulation effects.

In a second step, to test for domain-specific effects of FPC-TPJ versus FPC-OC tACS, we ran a GLMM where, in addition to the above-mentioned predictors, we added predictors for Task (0 = perceptual, 1 = social) and Experiment (0 = FPC-OC, 1 = FPC-TPJ) as well as all interaction terms with tACS, Accuracy and Difficulty (both as fixed and random effects), and a fixed-effects predictor for previous tACS condition. For the analysis of metacognitive efficiency, we computed individual Mratio values using the *metaSDT*-package (62) separately for each task and tACS condition with the SSE-method. We ran a GLMM regressing Mratio on dummy-coded fixed-effects predictors for tACS, Accuracy, Task, and Experiment as well as all interaction terms as random slopes in addition to participant-specific random intercepts. As a further measure of metacognition, we computed metacognitive noise, which quantifies the internal variability arising in confidence formation (45), typically manifesting as reduced metacognitive sensitivity. To this end, we followed the computational approach by Rahnev (63) to compute separate metacognitive noise values for each participant, tACS condition, and experiment (see Supplementary Materials Note S1 for more information on the model). We compared these values using the glmmTMB-package (75) to compute a GLMM with a Gamma distribution and log link function to account for the skewness of the parameter distribution. Here, we ran three separate models and regressed metacognitive noise on a dummy-coded fixed-effects predictor for tACS condition, Task (0 = perceptual, 1 = social), and Experiment (0 = FPC-OC, 1 = FPC-TPJ), as well as all interaction terms, including subject-specific random intercepts. In each model, we only included two tACS conditions, thus directly comparing in-phase versus sham, anti-phase versus sham, as well as in-phase versus anti-phase, respectively. In addition to this, we ran separate models, where we regressed metacognitive noise on a fixed-effects predictor for tACS condition (1 = in-phase versus sham, 2 = anti-phase versus sham), including subject-specific random intercepts, separately for tasks and experiments.

## Acknowledgements

AS was supported by an Emmy Noether fellowship (SO 1636/2-1) and a Heisenberg fellowship (SO 1636/3-1) from the German Research Foundation.

## Data Availability Statement

Behavioural data and analysis code are available on OSF (https://osf.io/ye8v3/overview) and are publicly available. Processed fMRI data are available upon reasonable request.

